# Glucocorticoid involvement in reproductive biology

**DOI:** 10.1101/2022.11.03.515013

**Authors:** Linda J. Mullins, Steven D. Morley, Christopher J. Kenyon, John J. Mullins

## Abstract

Oestrogen and progesterone play essential roles in the release of mature oocytes, the priming and cycling of the uterine lining, and the maintenance of mammalian pregnancy. Progesterone is synthesized *de novo* at the embryo implantation site in the mouse, during decidualization of the endometrium. During early stages of pregnancy, the locally produced progesterone is thought to act as an immunosuppressant, preventing rejection of the fetal allograft at the fetal-maternal interface. However, both uterine natural killer cells and dendritic cells express glucocorticoid receptor rather than progesterone receptor. The importance of glucocorticoids in early pregnancy is inferred from the presence of steroid receptors and the 11β-hydroxysteroid dehydrogenase enzymes, which modulate corticosterone action in the decidua, the trophoblast, the placenta, and the fetus. 11β-hydroxylase is the last enzyme in the metabolism of cholesterol to corticosterone and, in a mouse model of 11β-hydroxylase deficiency, complications of reproduction suggested its requirement for normal ovulation and uterine cell turnover. We present evidence that, in this model, folliculogenesis occurs normally but ovulation is inhibited, and abnormal uterine cell turnover ultimately leads to adenomyosis. Ovaries respond to a superovulation protocol by releasing oocytes and forming corpora lutea, and homozygous null blastocysts are capable of implantation, but the pregnancy is not maintained. We show that glucocorticoid is produced locally at the implantation site in control animals, revealing wide involvement of glucocorticoids in reproductive biology.

## Introduction

The involvement of oestrogen and progesterone in reproductive biology are widely acknowledged, however the evidence for glucocorticoid involvement during ovulation and pregnancy is largely circumstantial gleaned from the effects of synthetic glucocorticoids administered before and during pregnancy, the effects of maternal glucocorticoid levels induced by stress/poor diet and the results of impaired fetal/maternal glucocorticoid metabolism on resultant progeny (1). Intra-uterine growth restriction and fetal programming are recognised outcomes resulting from maternal stress during late pregnancy (2), while the presence of 11β-hydroxysteroid dehydrogenase enzymes in the uterus (3), the placenta (4) and the fetus (5) suggest the need to control fetal exposure to glucocorticoids. Indeed, in rodent models of Hsd11b2 knockout, progeny are found to develop hypertension from 5 weeks of age (6), partly as a result of exposure to high levels of glucocorticoids *in utero* (7). Glucocorticoids administered early in pregnancy may significantly increase pregnancy rate for IVF patients, though a similar beneficial effect on intra-cytoplasmic sperm injection (ICSI) is not evident (8, 9). Though high cortisol levels immediately post conception are associated with increased risk of miscarriage, and drugs that non-specifically block progesterone and glucocorticoid action are used to induce abortion (10), glucocorticoids are administered early in pregnancy to prevent female genital virilisation in patients with congenital adrenal hyperplasias (11).

Implantation of an embryo is a huge immunological challenge to the mother, yet the embryo is allowed to attach, and trophoblast cells are tolerated as they invaginate into the decidualizing tissue. This comes about through suppression of the innate maternal immune system - a complex process conventionally attributed to progesterone (12). Progesterone is produced by the corpus luteum, formed in the ovary once the oocyte has been released from the developing follicle. Key steroidogenic enzymes, cholesterol side-chain cleavage cytochrome P450 (P450scc) and 3β-hydroxysteroid dehydrogenase type VI (3βHsdVI), together with accessory proteins, are also expressed at the site of implantation (13). Thus, progesterone levels are increased in the decidua. By mid-gestation expression of steroidogenic enzymes transfers to the giant trophoblast cells, and then declines during the second half of pregnancy.

Uterine natural killer (uNK) cells (the predominant leukocyte population present in the endometrium at the time of implantation (14)) and uterine dendritic cells (uDC; potent antigen presenting cells) are clearly involved in the process of fetal-maternal tolerance (15). Progesterone appears to inhibit mature DCs and causes activated T-cells to produce LIF (essential for embryo implantation), macrophage colony stimulating factor (important for pregnancy development) and progesterone-induced blocking factor, PIBF. PIBF induces a predominantly Th2 environment and the Th2-type cytokines, including IL-4, block NK activity and macrophage functions. Recent in vitro evidence suggests that uDC and uNK cells not only ‘talk’ to each other, but are also modulated by trophoblast cells (reviewed (16)). Intriguingly, uNK cells express glucocorticoid receptors but not progesterone receptors (17, 18).

Previously we generated a mouse model in which 11β-hydroxylase, the final enzyme in cholesterol metabolism to glucocorticoid, was insertionally inactivated(19). Homozygote animals were initially extremely rare, indicating embryonic lethality. A surviving homozygote male proved to be viable and fertile, producing both male and female null progeny, which exhibited classic symptoms of CAH. Unexpectedly, null females were found to be infertile. Ovaries of Cyp11b1 null females contained pre-ovulatory follicles, but were completely devoid of recognisable corpora lutea, presumably due to high levels of circulating progesterone caused by the block in adrenal glucocorticoid synthesis. Instead, they contained lobular, amorphous, non-secretory, non-proliferative leutenized granulosa cells. Chronic exposure to high levels of progesterone also caused endometrial hyperplasia and adenomyosis in older null females. Continued investigation into the causes of infertility, has now led to the discovery that ovaries are responsive to a superovulation protocol, the uterus exhibits increased cell division /turnover and glucocorticoids are actively synthesised at the site of implantation.

## Results

### Cell proliferation in ovaries and uterus of null females

Having previously shown that ovaries of null females were devoid of corpora lutea, negative for proliferating cell nuclear antigen (PCNA) and were not actively proliferating (as assessed by BrdU turnover within 2 hours of sacrifice (19)), we conducted a pulse-chase experiment, where BrdU was administered for one week, and then tissues were harvested 6 weeks later. WT animals retained punctate label in granulosa cells, suggesting that cells of some primary, secondary, and large pre-antral follicles, but not primordial follicles had undergone cell division during the chase period. Large quantities of label were retained in nuclei of stromal cells surrounding the corpus luteum of WT females, suggesting quiescence (Fig.1a, c). Despite lacking corpora lutea, Cyp11b1 nulls showed normal folliculogenesis, and BrdU staining was found in broadly similar patterns (Fig.1b, d).

**Figure 1:**
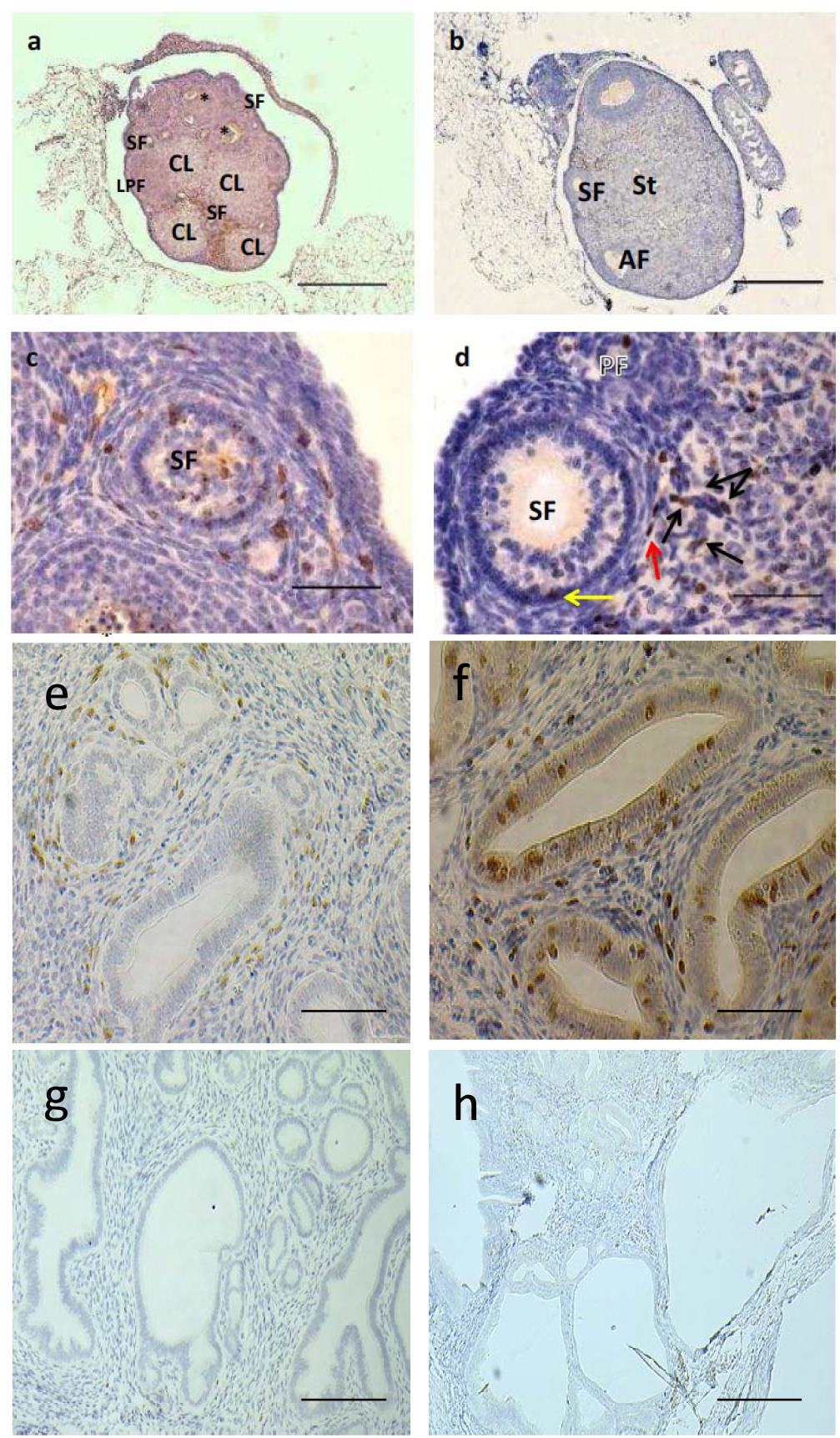
BrdU distribution following pulse and six-week chase in (a, c) WT ovary; (b, d,) Cyp11b1-/- ovary; wild type uterus (f) Cyp11b1 KO uterus. (g,h) morphology of WT and Cyp11b1 KO uterus. Label-retaining cells in ovary are; granulocytes (yellow arrow); theca cells (red arrow); stroma (black arrows). CL - corpus luteum; LPF – large preantral follicle; SF – secondary follicle; AF – atretic follicle; St – stroma.

The uteri were also analysed following the BrdU pulse-chase experiment. Many more label-retaining cells were found in the Cyp11b1 null uterus than in WT, suggesting a more protracted turnover of putative stem cells (Compare Fig.1f with Fig.1e). Also, Ki67 double labelling suggested a much higher level of active cell division in the null female uterus, reflecting the previously observed hyperplasia (data not shown).

### Pregnancy following superovulation

Previously, we determined that the ovaries of null females responded to superovulation, releasing normal (15-20) numbers of oocytes. To see if null females were equally capable of becoming pregnant and sustaining pregnancy, 5 null females were superovulated, mated with Cyp11b1-null males, plug checked and then culled at e6.5 or e7.5. The ovaries were found to contain multiple corpora lutea, when visualised by optical projection tomography (OPT; Fig.2a) with over 20 CLs counted through the 3D stack (Supplementary video S1). By immunohistochemistry, the CLs stained positive for pdcd4 (Fig.2b) and looked indistinguishable from WT (not shown). However, staining with Col4a2 was considerably reduced in the corpus luteum of the Cyp11b1 KO ovary (Fig.2e and f) compared to WT (Fig2c and d). Only two null females became pregnant, carrying 5 and 2 implantation sites respectively. The implantation sites were scanned by OPT, frozen for RT-PCR analysis or processed for immunohistochemistry.

**Figure 2:**
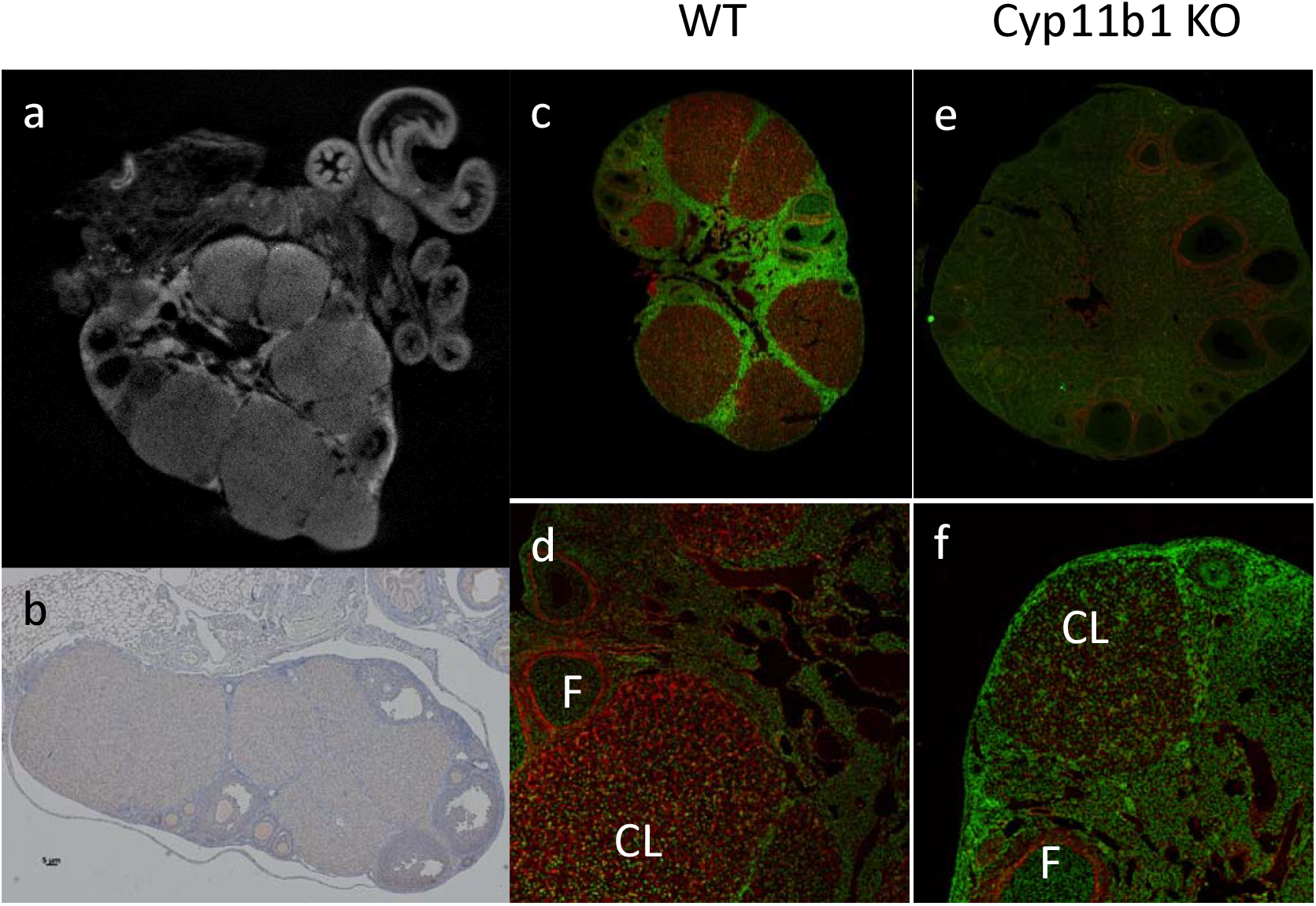
(a) corpus lutea in Cyp11b1-/- ovary following superovulation, revealed by OPT; (b) Cyp11b1-/- ovary section immunostained with anti-pdcd4; (c, d) ovary from WT female or (e, f) superovulated Cyp11b1 KO female, stained with anti Col4a2; (e, f) higher magnification of Col4a2 stained follicle (F) and corpus luteum (CL)F

### Optical Projection Tomography analysis

OPT analysis allowed us to measure accurately the uterine diameter at the implantation sites (Fig.3a-d) (and Supplementary videos S2 to S4). Implantation sites at both e6.5 and e7.5 were significantly smaller (2.365 nm and 1.708nm respectively), than time-matched implantation sites from heterozygous females (e6.5 – 3.176nm). Interestingly, OPT allowed us to reconstruct and visualise the internal structure of the implantation site. Comparison with images from equivalent Theiler stages in the mouse atlas (http://www.emouseatlas.org/emap/ema/home.html) suggested that the development of the homozygous embryos in homozygous females was either delayed, decidualization was impaired or pup resorption was underway.

**Figure 3:**
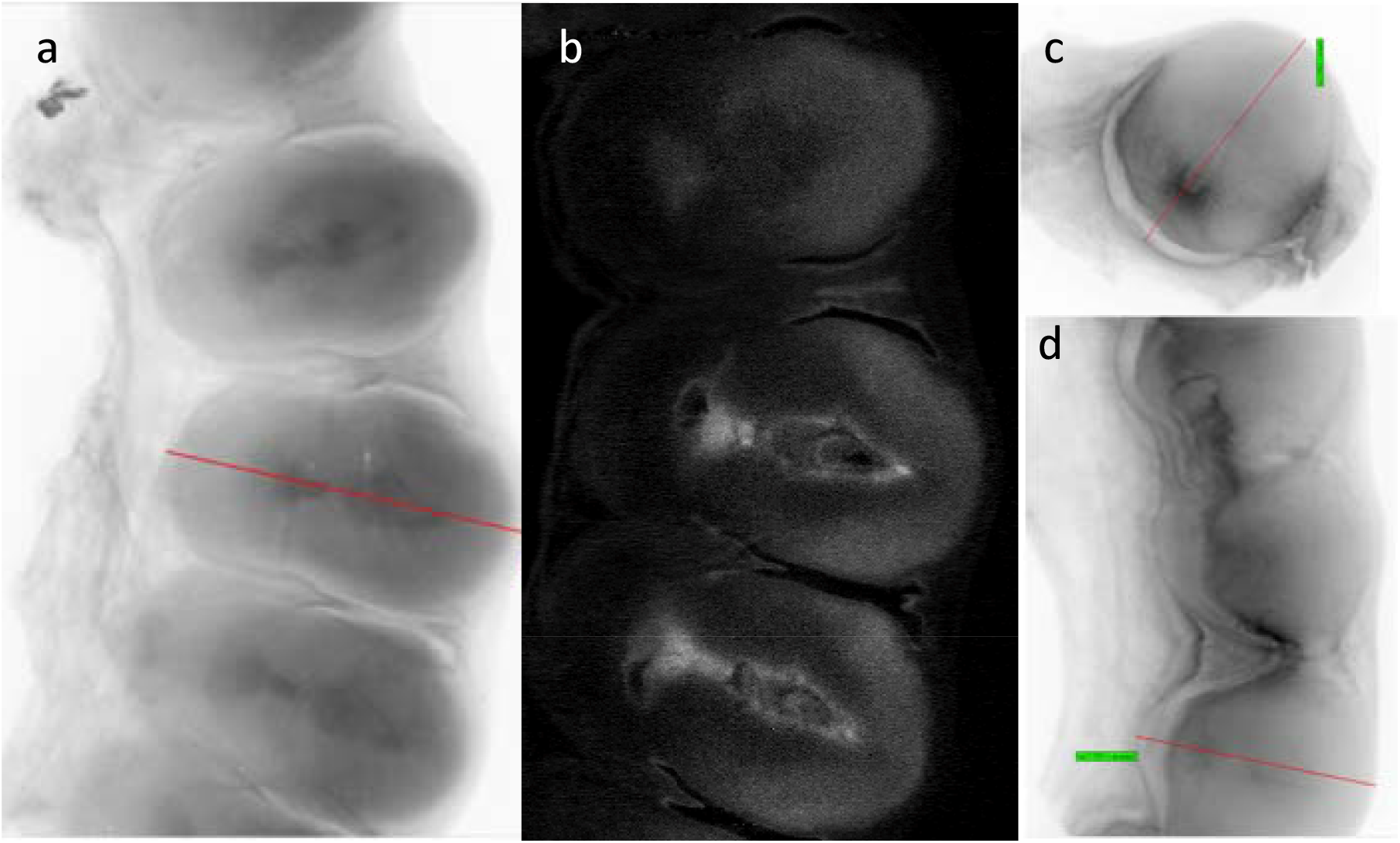
OPT scan of WT e6.5 implantation sites (sagittal section) using (a) brightfield and (b) autofluorescence. Brightfield OPT scans of (c) e6.5 and (d) e7.5 Cyp11b1 KO implantation sites

### In situ hybridization

To check for the presence of Cyp11b1 mRNA, *in situ* hybridization of implantation site sections was carried out, using sense and antisense probes to both Cyp11b1 and the closely related gene Cyp11b2 (aldosterone synthase). Hybridization, using the Cyp11b1 antisense probe, was observed in WT (Fig4a-c) and heterozygous but not nullizygous implantation sites. There was no hybridization using either the Cyp11b1 sense probe or sense or anti-sense cyp11b2 probes in implantation samples from any genotype.

**Figure 4.**
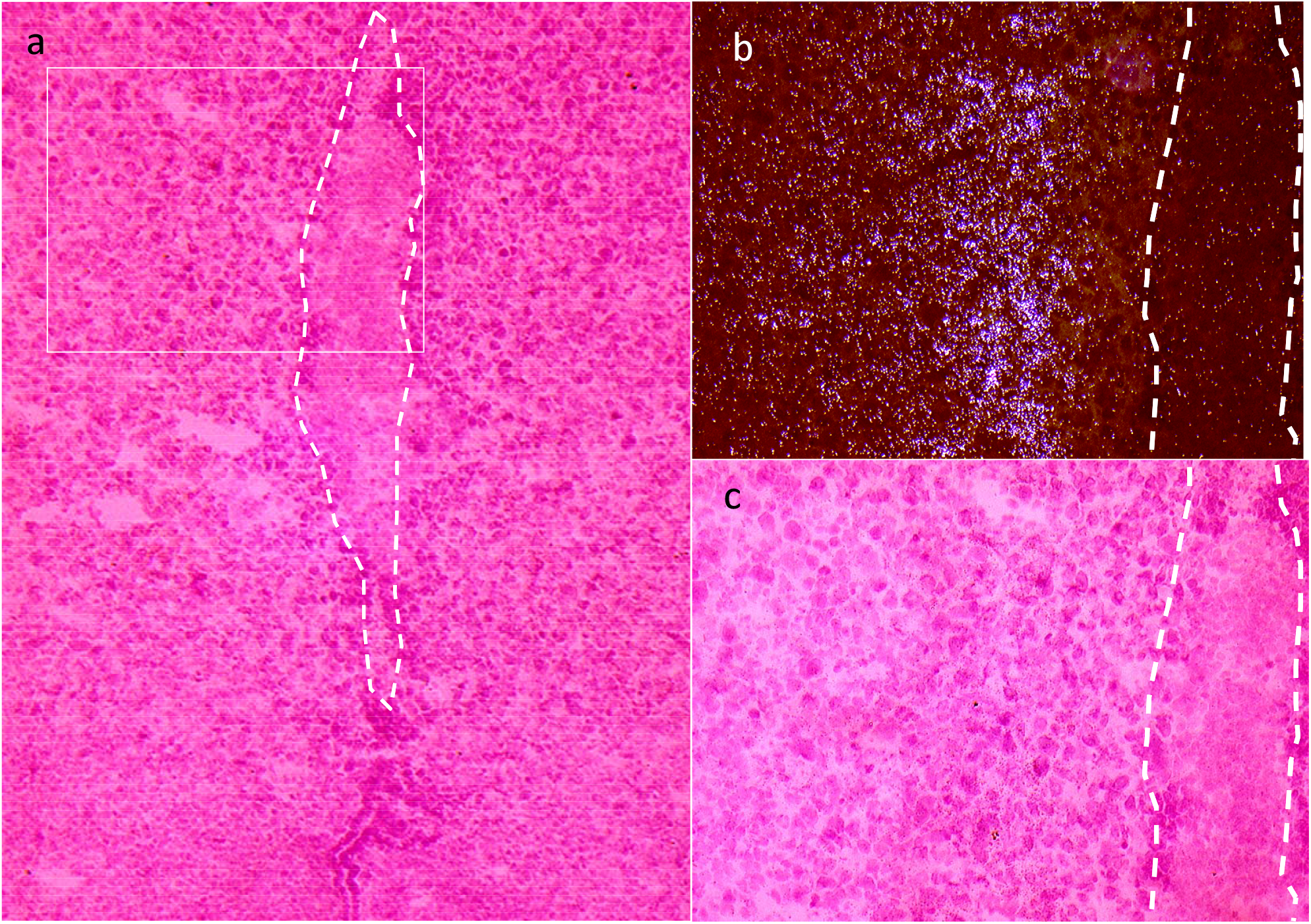
(a, c) Embryo location (e7.5 WT) visualized with H&E stain and marked with dotted line. (b) in situ of implantation site showing Cyp11b1-speciific staining in decidual tissue flanking the embryo-derived tissue.

### Reporter fluorescence/immunofluorescence

Fluorescence of the ECFP reporter, which was incorporated during targeted knock-out of the *Cyp11b1* gene was observed in the zona fasciculata of the adrenal gland (Fig.S1a-d) and additionally, the uterus of knockout females (Fig.S1e), using immunofluorescence antibody against GFP. Staining was observed in decidual tissue and was stronger at the pole.

### Immunohistochemical analysis of implantation sites

WT implantation sites were screened by immunohistochemistry, using anti-glucocorticoid receptor (GR) antibody. GR was found throughout the decidua, and was largely nuclear, but there was some cytoplasmic staining in a subset of cells (Fig.S2a). Nuclear GR stain was also seen at the implantation site of a homozygous female (Fig.S2b).

It is well known that progesterone is synthesized at the implantation site. WT implantation sites were assessed for the presence of 21-hydroxylase (using appropriate antibody), which was found to stain weakly in maternal decidual tissue at e7.5 implantation sites (Fig.S2c-d). Unfortunately, equivalent assessment of Cyp11b1 expression was not possible because of adverse background staining using anti-11BOH antibody or anti-GFP antibody for nullizygous samples. Pancytokeratin, a marker of extraembryonic trophoblasts was indistinguishable between WT and Cyp11b1 KO implantation sites using a pancytokeratin antibody (Fig.S2e-f). Infiltration of immune cells at the implantation site was assessed using F4/80 antibody, but again was indistinguishable between WT and Cyp11b1 KO implantation sites (Fig.S2g-h).

### RT-PCR analysis

Implantation sites and inter-implantation uterine sections (where these were present) were carefully dissected from e5.5 to e8.5 pregnant WT mice, and analysed by RT-PCR, using primers for *Cyp11b1, Cyp21* and *Hsd11b2. Cyp11b1* mRNA was expressed specifically in the implantation sites from e5.5 onwards but was barely detectable in inter-implantation tissues (Fig.5a) or in uterine tissue of plugged females at e4.5 (data not shown). *Cyp21* mRNA was detected at a low level in all tissues, and at all time points assessed (Fig5b), while *Hsd11b2* mRNA expression within the implantation sites was lower than inter-implantation expression (Fig5c).

**Figure 5:**
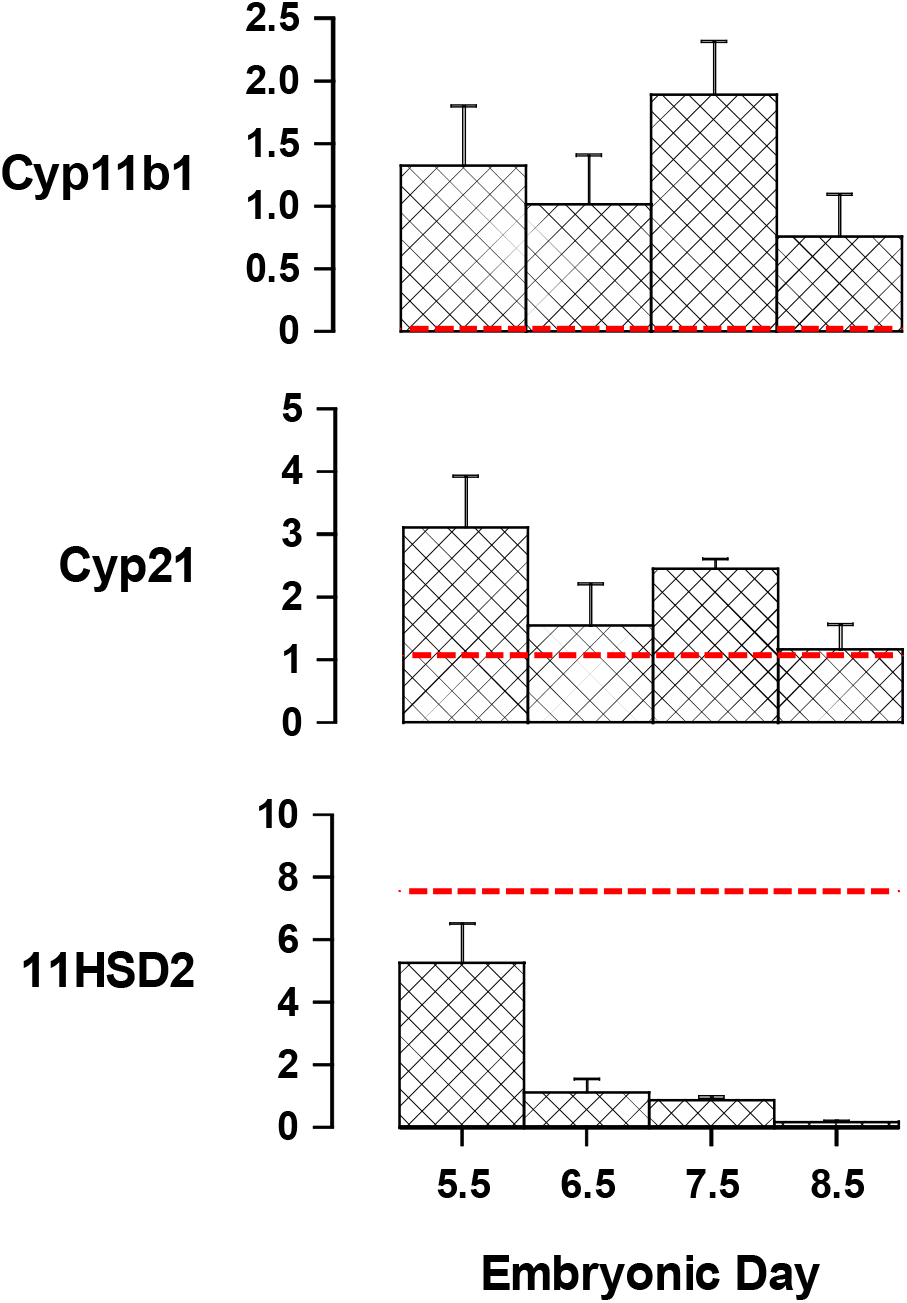
(a-c) Graphs showing RT-PCR analysis of implantation sites from e5.5 to e8.5 for Cyp11b1, 21-OH and Hsd11b2. In each case, the red line indicates the level of expression seen in inter-implantation site tissue at e5.5

**Figure 5:**
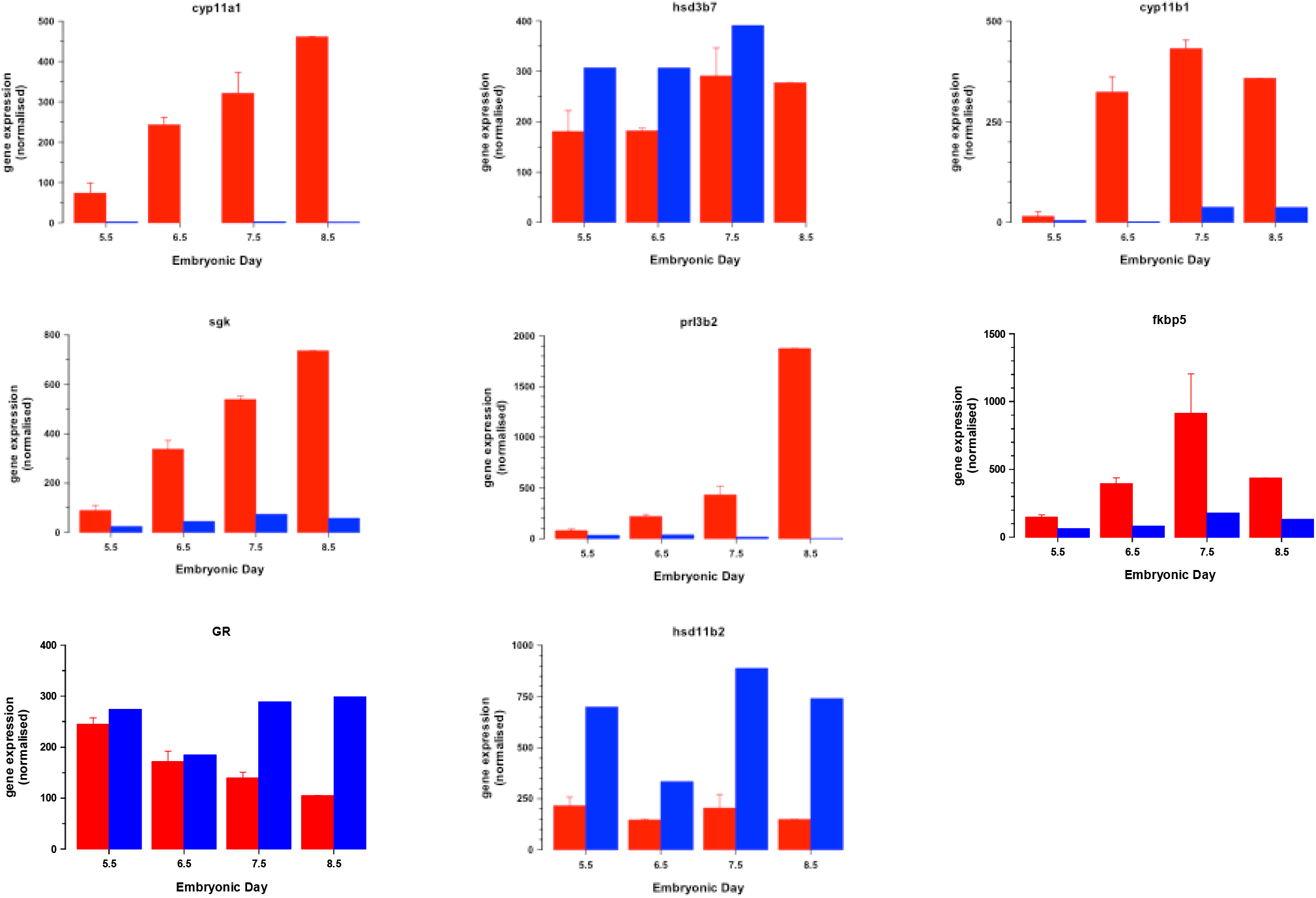
(d-k) RT-PCR analyses of mRNAs, encoding steroidogenic enzymes of the cholesterol metabolic pathway (Hsd3b7 and Cyp11a1), glucocorticoid-regulated genes (sgk1, Prl3b2 and Fkbb5) and glucocorticoid responsive or metabolising genes (GR, Hsd11b2). Red bars – implantation sites; blue bars – inter-implantation site tissue

Analysis was extended to confirm the presence of mRNA encoding other steroidogenic enzymes of the cholesterol metabolic pathway (*Hsd3b7* and *Cyp11a1*), glucocorticoid-regulated genes (*sgk1, Prl3b2* and *Fkbb5*) and glucocorticoid signalling genes (*Nr3c1* (GR), *Hsd11b2*) (Fig.5d-k). Similar patterns of gene expression were observed using tissues isolated from super-ovulated pregnant WT females indicating that superovulation does not qualitatively affect the expression of any of these genes (data not shown). *Cyp11b1* mRNA was not detectable in implantation sites from the Cyp11b1 null females carrying homozygous implants (data not shown).

## Discussion

Infertility in the Cyp11b1-null female was presumed to result from the increased progesterone levels caused by the block in glucocorticoid synthesis, rendering the female ‘pseudopregnant’ and therefore anovulatory. Folliculogenesis appeared to be normal, and the turnover of putative stem cells, assessed by BrdU pulse chase, paralleled that observed in WT females. It has been shown that knockout of the glucocorticoid receptor in zebrafish results in impaired maturation capacity of follicles, leading to reduced ovulation (20). Additionally, it has been reported recently, that glucocorticoids play an important role in primate periovulatory follicle competency (21), since an increase in the corticol:corticosterone ratio of follicular fluid was associated with oocytes which went on to form competent blastocysts in vitro. These observations suggest that glucocorticoids may play a key role in oocyte maturation.

The effect of chronic exposure of the uterus to progesterone appeared to be a protracted turnover of putative stem cells and a much higher level of active cell division in the nullizygous uterus. This reflects the observed hyperplasia and ultimate resolution to adenomyosis. The failure of an oestrus cycle probably prevents the cyclical proliferation and apoptosis of uterine lining, leading to an accumulation of BrdU-stained cells. Recently, glucocorticoids have been shown to limit the regenerative capacity of endometrial stem cells (22)

Superovulation was found to overcome the progesterone block on oocyte release and at e6.5, several corpora lutea were seen in the ovary. This suggests that egg release, wound repair and corpus luteal hormonal function - priming the uterus for embryo implantation - continue once the block is removed. Similar observations were made in a C/EBPb knock-out model (23). The abnormal collagen staining observed in Cyp11b1 null superovulated ovary suggests that it may not be functioning normally, however.

Clear evidence of embryo implantation in Cyp11b1-null females was observed at e6.5 and e7.5. This suggested that implantation-competent blastocysts had successfully interacted with a receptive uterus, and that blastocyst activation, attachment and penetration were all achieved, together with endometrial decidualization. However, the number of implantation sites was significantly lower than the number of eggs released from the ovary (as measured by the number of copora lutea observed by OPT), and the size of each implantation site was smaller than time-matched heterozygous or WT implantation sites, suggesting that some part of the implantation/pregnancy maintenance program was compromised.

The pronounced paucity of homozygous knockout progeny derived from heterozygous mothers in this line may reflect a general deficiency in implantation competence of Cyp11b1 knockout embryos. When WT males were mated with SO nullizygous females, up to 20 obligate heterozygous implantation sites were counted. Our observations suggest that nullizygous embryos have a greatly reduced capacity to implant in nullizygous females and fail to develop beyond e7.5.

The involvement of glucocorticoids during pregnancy has previously been inferred, as discussed in the introduction. No *Cyp11b1* mRNA was expressed at e4.5 or in inter-implantation sites. We demonstrated the up-regulation of *Cyp11b1* mRNA at the implantation site, and by *in situ* hybridisation it was shown to be localised to decidual tissue surrounding the embryo. Together with increased levels of Hsd11b2 mRNA in inter-implantation tissue, our observations strongly suggest that corticosterone is both required at the implantation site and contained within it, in a temporal and spatial manner. The low level of 21-hydroxylase (demonstrated both at mRNA level and by immunohistochemistry) might effectively limit the proportion of progesterone converted to glucocorticoid at the implantation site.

It should be noted that glucocorticoid is not completely absent in the null animals since corticosterone is produced as an intermediate metabolite by aldosterone synthase (e.g. plasma levels are 10% that of WT). Also Hsd11b1, which converts inactive 11-dehydro to active cortisol/corticosterone, is increased by progesterone in human and ovine decidualizing stromal cells (24, 25) and also in mouse decidua (3).

After mining data from a microarray study comparing decidual and deciduomal tissue (26), a number of genes were chosen to investigate correlation of expression of genes regulated by glucocorticoid (*Sgk1, Prl3b2* and *Fkbb5*) or involved in glucocorticoid action or metabolism (*Nr3c1* (GR), *Hsd11b2*). All the glucocorticoid-responsive genes demonstrated clear correlation with *Cyp11b1* mRNA expression from e5.5 to e8.5. Of these, Sgk1 is of particular interest, since it has been shown to be essential prior to implantation, through its down-regulation of *Nedd4-2* and activation of the epithelial sodium transporter, ENaC (*Scnn1a*) in the luminal epithelium, which reduces luminal fluid and facilitates closure of the uterine cavity. During the window of receptivity, Sgk1 levels transiently decline (27, 28) and this coincides with the observed lack of *Cyp11b1* mRNA (e4.5). The expression of *Cyp11b1* mRNA was negatively correlated with the glucocorticoid receptor mRNA from e5.5 to e8.5. This may again reflect the need for tight control of glucocorticoid action at the site of implantation.

A possible role for glucocorticoids in early pregnancy is the moderation of pro-inflammatory cytokine interactions with prostaglandins (an anti-inflammatory effect) (16, 17) leading to fetal tolerance. Since uNK cells do not express progesterone receptors, any effect of progesterone must be indirect (18). Glucocorticoids may directly influence the cytokine profile of T cells and increase IL-4 production. Recently, it was shown that mifepristone (RU468), which is an antagonist of both progesterone and glucocorticoid, increases the cytotoxicity of human uNK cells by increasing perforin and ERK1/2 phosphorylation. Importantly, both effects were attenuated by cortisol but not progesterone (29). Additionally, it was shown that impaired corticosteroid signalling in decidualising stromal cells leads to increased uNK cell density(30).

In the absence of synthesized glucocorticoid – as with nullizygous fetuses in a nullizygous mother - implantation occurs, but the embryo may either be recognised as poor quality or fail to trigger some response in the decidual tissue, which may ultimately result in fetal reabsorption through breakdown of fetal-maternal tolerance. It was reported recently that multiple and different glucocorticoid receptor isoforms are produced by oocytes and blastocysts (31). This may lead to subtle alterations in the interaction of blastocysts at the site of implantation. Though we failed to demonstrate an increase in the presence of immune cells (via anti-F4.80 immunostaining) at the nullizygous implantation sites, it is tempting to speculate that corticosteroid specifically synthesised at the site of implantation, is essential for control of uNK number and cytotoxicity, and it is likely that fetal-maternal tolerance in the null implantation site would be compromised.

In conclusion we have demonstrated evidence to suggest a clear involvement of glucocorticoids during ovulation, uterine cell turnover during the oestrus cycle, and blastocyst maintenance following implantation, possibly through immune suppression. All three observations deserve further investigation.

## Materials and Methods

### Animal studies

All experiments were approved by the local ethics committee and conducted in accordance with UK Home Office regulations and the Animals (Scientific Procedures) Act 1986.

Animals were housed in standard cages, maintained on a 12 h light/dark cycle (7.00am to 7.00pm), and given free access to water and standard rat chow.

Superovulation was achieved by administration of pregnant mare’s serum gonadotrophin (Intervet; 20iu) at 9a.m., and human chorionic gonadotrophin (‘Chorulon’ Intervet; 30iu) 48 h later. They were then mated that evening and were plug-checked the following morning (e0.5).

### BrdU pulse chase experiment

Mice were infused with BrdU solution () by surgical implantation of a 0.1ml Alzet mini-pump (model 1007D. Palo Alto, CA, USA) in 10 month old WT or Cyp11b2 null mice for one week. The minipump was surgically removed and the animals were killed six weeks later. Tissues were fixed for 24 h in 4% neutral buffered formalin, transferred to 70% ethanol and embedded in paraffin wax.

### Optical Projection Tomography

Ovaries or implantation sites, which had previously been fixed in 4% paraformaldehyde, were rinsed briefly in PBS. They were then set within 1% low melting point agarose, mounted, dehydrated in methanol (24-48 h), and clarified with BABB (2: 1 benzoyl benzoate: benzyl alcohol; minimum of 4 hours). They were then scanned by Optical protection tomography (OPT) (32) using a Bioptonics 3001 OPT tomograph, either using brightfield or UV illumination (425 nm excitation filter with 40 nm band pass; 475 nm long pass emission filter; 1.048 Mpixel scanning resolution). Raw data (400 projections per scan at 0.9° increments) was subject to Hamming-filtered back projection using NRecon software (Skyscan, Belgium).

### In situ hybridization

In situ hybridization was carried out as described previously(33). Briefly, frozen sections (10um) were post-fixed in 4% paraformaldehyde and then hybridized overnight with [^35^S]UTP-labelled cRNA probes transcribed from PCR products of Cyp11b1 and Cyp11b2. Sections were then treated with RNAse A, washed to 0.1xSSC stringency, dried and exposed to film. Control sections were hybridized with labelled sense cRNA probes.

### Tissue processing

Tissues were harvested into RNA later, frozen on dry ice in OCT, fixed in 4% PFA ON at 4°C, or fixed in formalin overnight, as necessary. For perfusion fixation, animals were anaesthetized under isoflurane, and the infra-renal aorta was cannulated to allow retrograde perfusion of 150 ml fresh 4% PFA in PBS, pH 7.4. Tissues were then removed, trimmed and placed in 4% PFA overnight at 4°C, followed by 70% ethanol. All fixed tissues were embedded in paraffin wax and sequential 5μ sections cut. Where possible, implantation sites were immediately identified under a compound microscope. Otherwise, every fourth section was stained with Haematoxylin and Eosin to determine sections spanning the implantation site.

### Fluorescence

Frozen sections (10μ) were cut and mounted on microscope slides prior to imaging under either brightfield, or confocal microscope, as necessary.

### Immunohistochemistry

Sections were de-waxed, hydrated and heated in Novocastra Bond Epitope Retrieval Solution 1 pH 6.0 (ERI, Leica), and target antigens were detected using a Leica BOND-MAX robot.

Primary antibodies were mouse monoclonal anti-11βhydroxylase (1:150) or mouse anti-GFP (Living colours; 1:200), mouse monoclonal anti-21-hydroxylase (1:750), all requiring pre-incubation with rodent block M prior to primary antibody staining; anti-glucocorticoid receptor (1:300), rabbit anti-bovine pan-cytokeratin (1:1000) and anti-F4/80 (1:300) with mouse on mouse polymer and DAB, or horse radish peroxidase-conjugated secondary antibodies, as appropriate. Some sections were counterstained with H and E.

### RT-PCR

Total RNA was extracted from tissues using trizol and treated with DNAse free (Invitrogen; manufacturers protocols). cDNA was prepared using Superscript II first strand synthesis kit (Invitrogen). Taqman gene-specific RT-PCR primer pairs were used with the ABI Prism 7000 (as follows: *Cyp11b1* – Mm01204952_m1; *Cyp21a* – Mm00487230_g1; *Hsd11b2* – Mm01251104_m1) and results were analysed using the ABI Prism 7000 software. RT-PCR assays using the Lightcycler (Roche) with Roche universal probes and primer pairs are specified (Supplementary Table S1). All measurements were carried out in triplicate and normalised to controls PP1a, Tbp and 18S.

### Statistical analyses

Data are presented as the mean +/-SEM. Variables were compared by performing two-way ANOVA using Prism6. Mean values were compared using the Student’s t-test. A P<0.05 was considered statistically significant.

## Acknowledgements

We wish to acknowledge Prof Alan McNeilly and Drs David Brownstein, Judy McNeilly and Kerry McInnes for helpful discussions; the technical assistance of Ms Nicola Wrobel, Ms Lynn Ramage and Mrs. Nina Kotelevtseva, and bioinformatics assistance of Dr. Donald Dunbar. A number of post graduate projects undertaken by Kai Stuckenschneider, Lesley Fowler, Tianshi Guo, Maria Elena Camacho Moll, Maryam Sadraie, and Kirsty Shearer, contributed in part to this manuscript.

## Supplementary data

Figure S1: Fluorescence observed in adrenal gland of (a, c) Cyp11B1KO and (b, d) wild type male. (e) fluorescence of uterine strip taken from Cyp11b1KO female. (f) schematic of endometrium showing En – Endometrium; EnGl – endometrial gland; LaPr – Lamina propria; Lu – lumen; My myometrium. (Taken from Atlas of mouse histology – uterus 20X).

Figure S2: Immunohistochemical analyses of implantation sites from WT (a, c, e, g,) and Cyp11b1 KO animals (b, d, f, h,) screened using (a, b) anti-glucocorticoid receptor (GR); (c, d) anti-21-hydroxylase; (e, f) anti-pan cytokeratin; (g, h) anti-F4/80.

Video S1: superovulated nullizygous ovary (OPT; Sagittal)

Video S2: E6.5 nullizygous implantation site (OPT; transverse)

Video S3: E7.5 nullizygous implantation site (OPT; transverse)

Video S4: E6.5 heterozygous implantation sites (OPT; Sagittal)

**Table S1:**
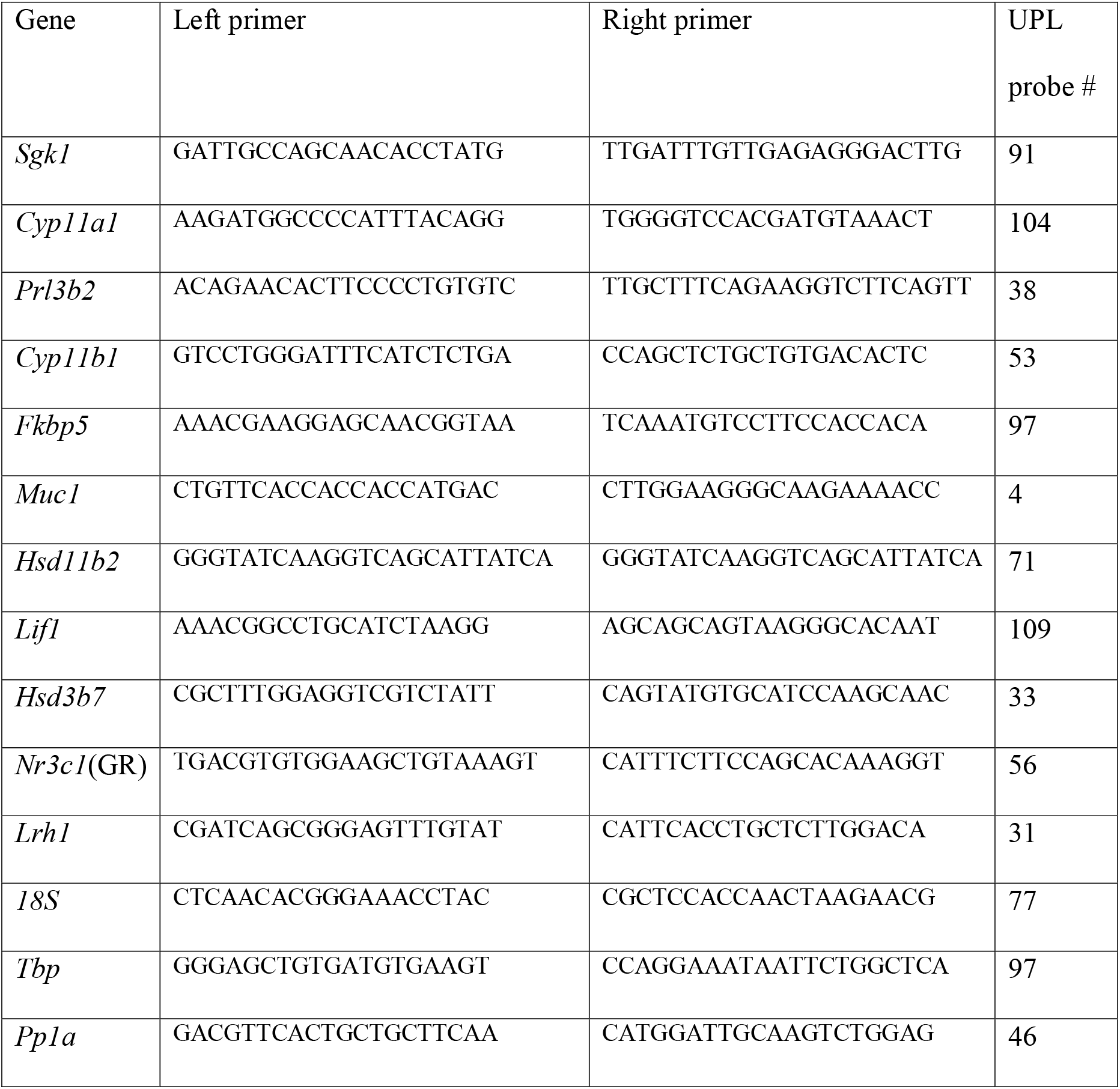
Primer pairs and UPL probes used for RT-PCR analysis using Roche Lightcycler

