## Supplemental figures S1 and S2 for "Glucocorticoid involvement in reproductive biology"

Supplementary Figure S1: Fluorescence observed in adrenal gland of (a, c) Cyp11B1KO and (b, d) wild type male. (e) fluorescence of uterine strip taken from Cyp11b1KO female. (f) schematic of endometrium showing En – Endometrium; EnGl – endometrial gland; LaPr – Lamina propria; Lu – lumen; My myometrium. (Taken from Atlas of mouse histology – uterus 20X).

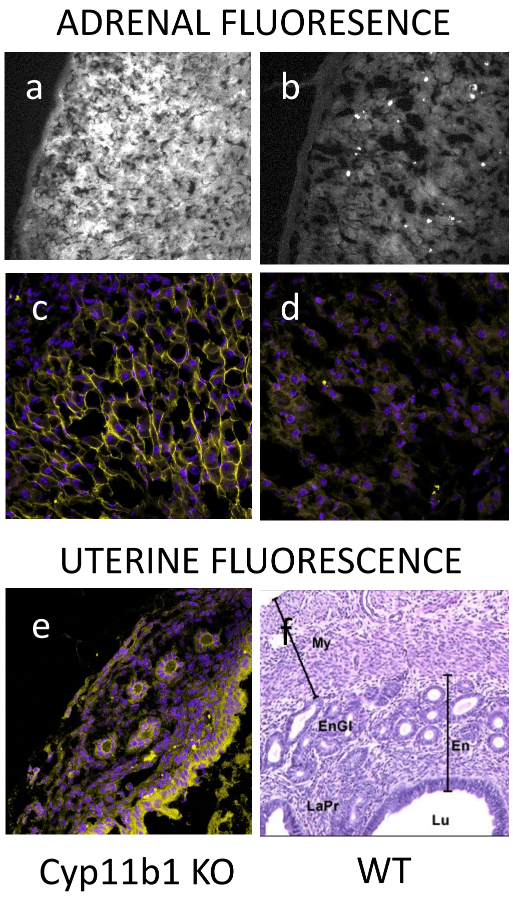

Figure S2: Immunohistochemical analyses of implantation sites from WT (a,c, e,g,) and Cyp11b1 KO animals (b, d, f, h,) screened using (a, b) anti-glucocorticoid receptor (GR); (c, d) anti- 21-hydroxylase; (e, f) anti-pan cytokeratin; (g, h) antiF4/80.

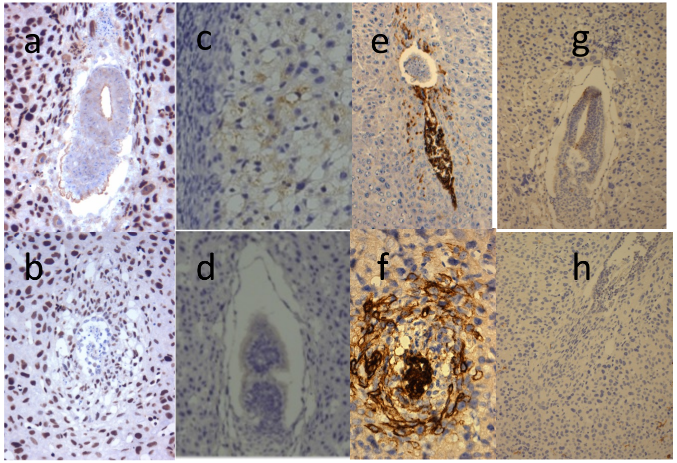
